# *Trypanosoma brucei* from pigs in sleeping sickness foci from Côte d’Ivoire is structured into clonal and strongly subdivided populations

**DOI:** 10.1101/2025.07.03.662103

**Authors:** Martial Kassi N’Djetchi, Thierry De Meeûs, Barkissa Mélika Traoré, Adeline Ségard, Sophie Ravel, Jacques Kaboré, Abe Allépo Innocent, Dramane Kaba, Djakaridja Berté, Thomas Konan, Bamoro Coulibaly, Pascal Grébaut, Geraldine Bossard, Bruno Bucheton, Mathurin Koffi, Vincent Jamonneau

## Abstract

Human African trypanosomiasis (HAT), or sleeping sickness, is currently targeted for elimination. The etiologic agent of HAT is a trypanosome belonging to the species *Trypanosoma brucei* (*Tb*) s.l., a unicellular parasite transmitted by tsetse flies. *Tb* s.l. consists of three subspecies: *Tb brucei* (*Tbb*), *Tb gambiense* (*Tbg*) and *Tb rhodesiense* (*Tbr*). These subspecies are morphologically indistinguishable and classified according to host in which they are found, type of disease and geographical distribution. During the last few decades, there has been considerable effort to genetically characterize *Tb* s.l. isolated from domestic and wild animals in order to better evaluate the impact of animal reservoirs on the epidemiology of HAT. To assess genetic diversity of *Tb* s.l. stocks circulating in three endemic or historical HAT foci in Côte d’Ivoire, we conducted a population genetics study of these parasites from animal hosts. Biological and isolated stock samples collected from pigs and reference stocks were tested with the primers of the *Trypanosoma gambiense-*specific-glycoprotein gene (TgsGP) and were genetically characterized with eighteen microsatellite loci. The few TgsGP positive samples found in pigs did not fit into *Tbg* reference stocks as regard to their microsatellite profile. These samples indeed were positioned far outside the lineage of this subspecies in the dendrogram built from genetic distances. We also found that in Ivoirian foci, *Tbb* populations (animal trypanosomes) were structured as several geographically strongly isolated units, localized in the vicinity of small villages. In these villages, data were compatible with a strict clonal propagation of *Tbb*. This is in variance with other published data on that subspecies. This study also confirmed the need to develop better tools to explore the relationships between *Tbb* and *Tbg*. Because sustainable elimination of HAT could be compromised, we also insist on the need to study the epidemiological role of potential animal and/or human reservoirs for *Tbg*.

## Introduction

The causative agent of human African trypanosomiasis (HAT), also known as sleeping sickness, is *Trypanosoma brucei* (*Tb*) s.l., a unicellular parasite transmitted by tsetse flies of the genus *Glossina*. *Trypanosoma brucei* infects human as well as a variety of domestic and wild animals in sub-Saharan Africa (Lejon et al., 2025).

*Trypanosoma brucei* s.l. is subdivided into three subspecies based on extrinsic criteria (host range, pathogenicity in human and geographical distribution). *Trypanosoma brucei gambiense* (*Tbg*) and *Tb rhodesiense* (*Tbr*) are both human-infective and responsible for a chronic form in West and Central Africa and an acute form in East Africa, respectively (Lejon et al., 2025). These taxa are also reported to infect animal reservoirs as mentioned in the literature: Lejon et al. (2025) for *Tbg* and Gibson (2002) for *Tbr*. *Trypanosoma brucei brucei* (*Tbb*), theoretically non-pathogenic to humans, is a parasite of domestic and wild animals causing animal African trypanosomosis (AAT or nagana) throughout the region delimited by the tsetse belt in sub-Saharan Africa (Hoare, 1972). With the advent of molecular methods, *Tbg* was subdivided into *Tbg* group 1 (*Tbg*1), a homogenous genetic entity responsible of most HAT cases (Gibson et al., 1986) and the rarest *Tbg* group 2 (*Tbg*2) recently defined as all human-infective *Tb* trypanosomes from West and Central Africa that do not fit into *Tbg*1 using molecular markers (Jamonneau et al., 2019).

With 747 and 799 cases reported in 2021 and 2022 respectively mainly in Democratic Republic of the Congo, Central African Republic and Angola, *gambiense* HAT (g-HAT) responsible for 94% of the total number of reported cases, is currently targeted for elimination of transmission (zero new cases during a period of five consecutive years) (Franco et al., 2024). However, the epidemiological role of domestic or wild animal, and/or human reservoirs, which could compromise this objective, is still under debate (Koffi et al. 2015; Büscher et al., 2018; Mehlitz et al., 2019).

In Côte d’Ivoire, important control efforts have led to a significant reduction of the prevalence of the disease (Dje et al., 2002; Kaba et al., 2006; Courtin et al., 2010; Koffi et al., 2016; Koné et al., 2020). Thanks to these efforts, the country has achieved the elimination of g-HAT as a public health problem in 2020 (Kaba et al., 2023), although a few cases were still diagnosed in the late 2010s (Koné et al., 2021). Potential animal reservoirs of *Tbg* have been described for a long time in HAT foci (Mehlitz,1986; Jamonneau et al., 2004). These animal reservoirs are now suspected to be partly responsible for the persistence of the disease (Kambire et al., 2012; Koffi et al., 2016; Kaba et al., 2023). Moreover, the existence of human reservoirs (asymptomatic and undiagnosed) cannot be ruled out (Koffi et al., 2009; Jamonneau et al., 2012). In recent studies conducted in the Bonon and Sinfra endemic HAT foci and the Vavoua historical focus, high prevalence of *Tb* s.l. was observed in domestic animals, mainly in pigs, due to a higher exposure of these animals to tsetse flies (N’Djetchi et al., 2017; Traoré et al., 2021). However, discordant results were obtained with the molecular and serological tools used. The specificity of the PCR targeting the *Trypanosoma gambiense*-specific-glycoprotein gene (TgsGP) described as a suitable method for the specific detection of *Tbg* (Gibson et al., 2010; Radwanska et al., 2002) was even questioned and both the existence and the epidemiological role of an animal reservoir for *Tbg* still remains unclear (N’Djetchi et al., 2017; Traoré et al., 2021).

The aim of the present study was to explore the population genetics of trypanosomes collected in the field from pigs from these three Ivorian gHAT foci (Bonon, Sinfra and Vavoua), and compare it to several reference stocks of *Tb* s.l. Our findings bring clarifications on the reproductive mode, dispersal and epidemiology of *Tbb*, as it also brings more confusion on the real specific or sub-specific status of the members of the entity named *T. brucei*.

## Material and Methods

### Ethical statement

Sample collection was conducted by a veterinarian of the Laboratoire National d’Appui au Développement Rural (Ministry of Agriculture) within the framework of epidemiological surveillance activities supervised by the HAT National Elimination Program (HAT NEP). Local authorities require no ethical statement. Any veterinarian may carry out blood sampling on domestic animals, with the authorization of the owner. No samples other than those for routine screening and diagnostic procedures were collected. Breeders gave their consent for animal sampling after explaining the objectives of the study. A deworming treatment (Bolumisol, Laprovet) was freely provided to all pigs sampled.

### Study areas

All samples included in this study were from Bonon and Sinfra endemic HAT foci and from the Vavoua historical HAT focus in the Western Center part of Côte d’Ivoire (Figure 1). These foci fall into the distribution area of tsetse flies (*Glossina*) (Laveissière et al., 1981; Ravel et al., 2007; Kaba et al., 2021). In these zones, AAT is mostly due to *Tbb* and *T. congolense* that mainly affect both pigs and cattle (N’Djetchi et al., 2017; Traoré et al., 2021).

**Figure 1:**
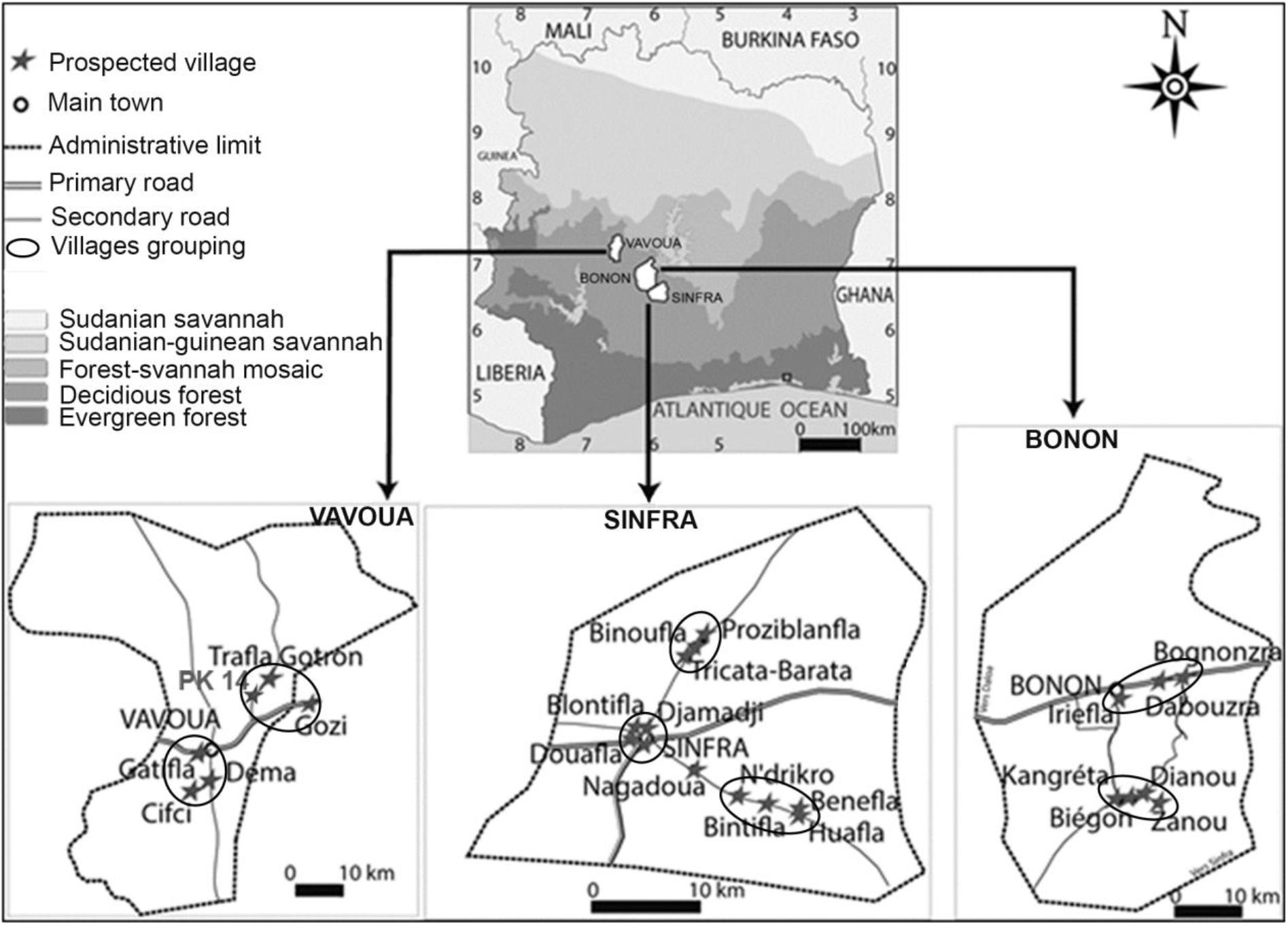
Study areas and sites of sampling of *Trypanosoma brucei* s.l. from pigs, indicating villages (black stars), village groupings (black ellipses, Nagadoua staying alone), and foci (Vavoua, Bonon and Sinfra). This figure was created by the mapping service of the “Unité de Recherche Trypanosomoses” based at Institut Pierre Richet (Bouake, Côte d’Ivoire).

Control efforts, conducted from 1992 until present, largely contained the HAT epidemic in Sinfra (1992-1997) and in Bonon (1998-2004) (Dje et al., 2002; Kaba et al., 2006; Kambire et al., 2012; Kaba et al, 2021). Nevertheless, few cases are still diagnosed each year (Kambire et al., 2012; Koffi et al., 2016; Koné et al., 2021; Kaba et al., 2023). The Vavoua HAT focus was epidemic in the 1970’s (Duvallet & Stanghellini, 1979) and thanks to control efforts, less than five HAT cases were reported yearly at the end of the 1990s (Dje et al., 2002). From the early 2000s, only four cases were still diagnosed (Simarro et al., 2010) and the last one was reported in 2011 (Traoré et al., 2021).

### Sampling

The number and the description of the selected field samples according to the focus, the year, the sampling method and the village are given in Table 1. Regarding the Vavoua focus (2017), we used 20 biological samples (BS) consisting of 500 μL of buffy coat obtained from 9 mL of heparinised blood collected from pigs and 20 isolated stock samples (IS) already described in Traoré et al.(2021). Both selected BS and IS, were positive with the TBR1-2 PCR (Moser et al., 1989) specific of *Tb* s.l. All BS were negative for the single round PCR targeting the TgsGP gene specific of *Tbg* (Radwanska et al., 2002). Five of the IS were TgsGP PCR positive: one giving a high intensity band and four a low intensity band (Traoré et al., 2021).

**Table 1:**
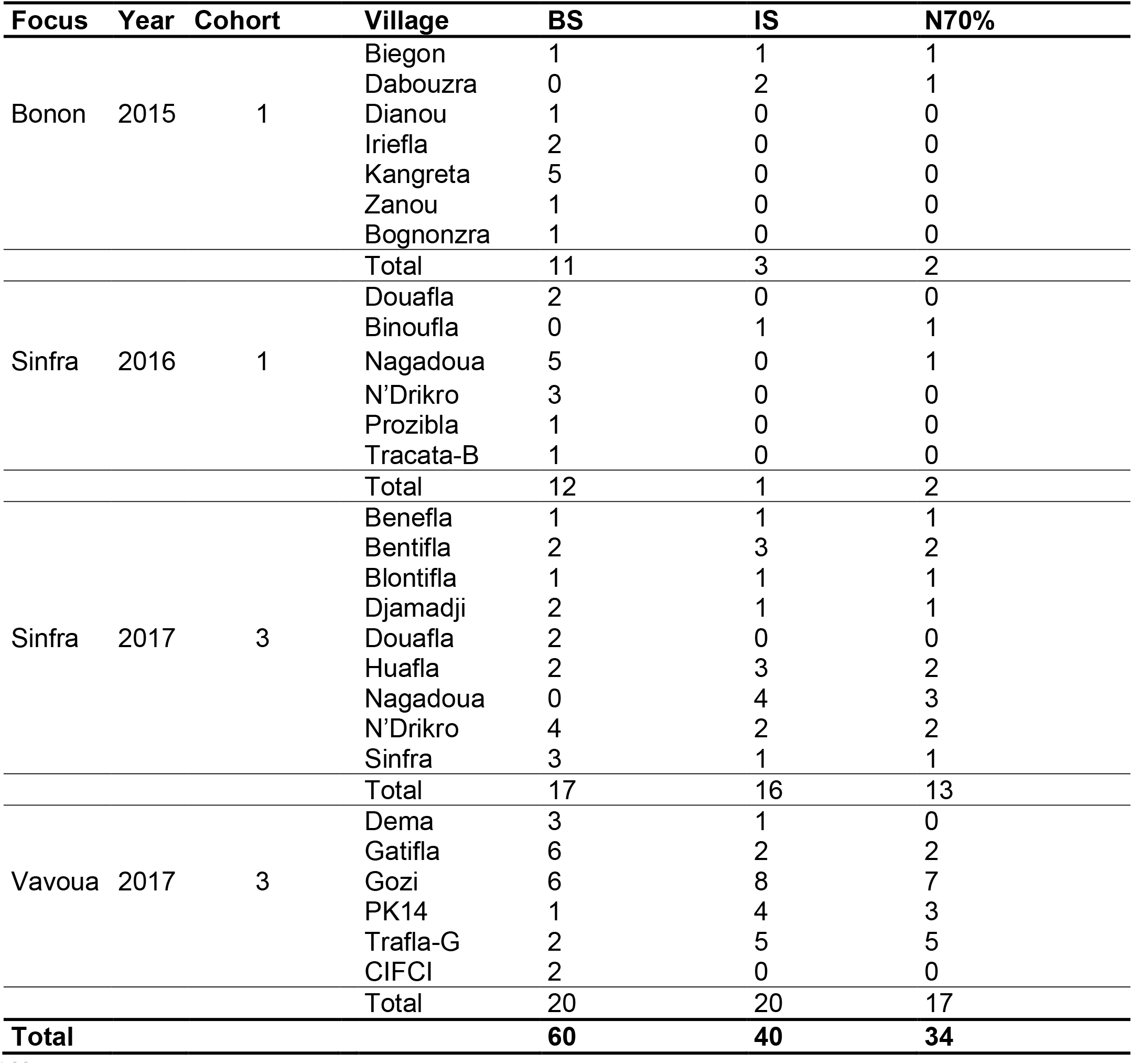
Description and number of samples used for the amplification of microsatellite loci of *Trypanosoma brucei* s.l. isolates from pigs in Côte d’Ivoire. Data are given regarding focus, year of sampling and cohort. We considered two different periods of six months as two different cohorts (Kaboré et al., 2011), the first corresponding to samples collected between December 2015 and May 2016 (cohort C1) and the second corresponding to samples collected between December 2016 and May 2017 (Cohort C3). The number of samples regarding the village, the sampling method (BS: biological samples, IS: isolated stock samples), as well as the number of samples that had at least 70% of interpretable loci used for the population genetics parameter estimates (N70%) are also indicated.

In Bonon in 2015 and in Sinfra in 2016 and 2017, the sample selection was also performed on BS (11 and 29, respectively) and IS (3 and 17, respectively) collected from pigs in the frame of studies on AAT epidemiology as previously described (Traoré et al., 2021). All these pigs tested positive for the trypanosome-detection buffy coat technique (Murray et al., 1977) and both BS and IS were positives to the TBR1-2 PCR but negative to the single round TgsGP PCR.

DNA from 27 reference stocks of *Tbb* (n=10)*, Tbg1* (n=13) and *Tbg2* (n=4) were also included in this study (Table 2).

**Table 2.** : Reference stocks of *Trypanosoma brucei sp* used for microsatellite loci characterisation, and taxonomic identification. Information provided for each stock includes the host species, geographical origin, year of isolation, and corresponding reference.

| Species | Stocks | Host | Origin | Year | References |
| --- | --- | --- | --- | --- | --- |
| *Tbg1* | B4F303 | Human | CI | 2004 | Koffi et al. (2007) |
|  | 107_4 | Human | CI | 1997 | Jamonneau et al. (2000) |
|  | B4U163 | Human | CI | 2004 | Koffi et al. (2007) |
|  | JUA | Human | Cam | 1979 | Stevens et al. (1992) |
|  | A005 | Human | Cam | 1988 | Stevens et al. (1992) |
|  | OK/ITMAP | Human | Con | 1974 | Stevens et al. (1992) |
|  | P26F | Pig | Cam | 1998 | Nkinin et al. (2002) |
|  | D12k | Sheep | Con | 1980 | Godfrey et al. (1990) |
|  | DIBBYEUG | Human | CI | 1998 | Jamonneau et al. (2000) |
|  | B4_K5 | Human | CI | 2004 | Koffi et al. (2009) |
|  | B4_Y221 | Human | CI | 2004 | Koffi et al. (2009) |
|  | B4C191 | Human | CI | 2004 | Kaboré et al. (2018) |
|  | T59_3 | Human | CI | 2002 | Koffi et al. (2009) |
| *Tbg2* | ABBA | Human | CI | 1983 | Hide et al. (1990) |
|  | MURAZ03 | Human | CI | 1979 | Godfrey et al. (1987) |
|  | TH2 | Human | CI | 1978 | Mehlitz et al. (1982) |
|  | FEOS | Human | Togo | 1961 | Tait et al. (1984) |
| *Tbb* | SW94_87 | Pig | DRC | 1987 | Truc et al. (1991) |
|  | DC8 | Pig | CI | 2001 | Jamonneau et al. (2004) |
|  | DC9 | Pig | CI | 2001 | Jamonneau et al. (2004) |
|  | EATRO1125 | Antilope | Ug | 1966 | Gibson et al. (1980) |
|  | P11F | Pig | Cam | 1998 | Simo et al. (2005) |
|  | P17F | Pig | Cam | 1998 | Nkinin et al. (2002) |
|  | GAOUA89 | Zebu | BF | 1989 | Simo et al. (2005) |
|  | STIB215 | Lion | Tan | 1971 | Gibson et al. (1980) |
|  | TSW65 | Pig | CI | 1982 | Stevens et al. (1992) |
|  | TSW53 | Pig | CI | 1982 | Stevens et al. (1992) |

### Microsatellite genotyping

Trypanosomes of the *gambiense* lineage are diploid (Cooper et al., 2008). These parasites can practice meiotic sex, as demonstrated between stocks of *Tbb* and *Tbg*2 (MacLeod et al., 2005), though at an undetectable rate (if any) (Koffi et al., 2009; Weir et al., 2016). DNA from BS was extracted from 500 μL of buffy coat using the DNeasy Blood and Tissue kit (Qiagen, Valencia, CA, USA) following manufacturer’s instructions. DNA from IS was extracted according to the same process, after having re-suspended the parasite pellets in 200 μL PBS.

All DNA samples were genotyped using four microsatellite primers (MisatG4, MisatG9, MicBg5 and MicBg6) previously designed by screening for dinucleotide repeats from a genomic bank of a *Tbb* strain (Koffi et al., 2007) and then labelled as *brucei* primers (BP). We also designed 14 new primers in the frame of this study. They were obtained from the genomic DNA of *Tbg* reference strain B4F303 (Koffi et al., 2007) and were labelled as *gambiense* primers (GP). These primers (name, repeated unit, sequence and annealing temperature) are described in the Supplementary file S1.

For BP, DNA samples were processed as previously described with a two-step amplification using a nested PCR (Kaboré et al., 2011). For GP, amplification conditions are described in the Supplementary file S1. After PCR amplification of microsatellite loci, alleles were routinely resolved on an ABI 3500 XL sequencer (Applied Biosystems, USA). This method allows multiplexing and the use of four dyes (6-FAM: violet-blue; PET: red, VIC: green; and NED: yellow). Allele calling was done using GeneMapper V 4.1 software (Applied Biosystems) and the size standard GS600LIZ short run. Amplification failures were noted in case of absence of amplification product or when the profile could not be interpreted.

### Data analyses

All data sets were designed for Create (Coombs et al., 2008) and converted into the appropriate format for the different analyses.

Comparisons of amplification success between BS and IS or between BP and GP were undertaken using all the individual samples (Tables 1 and 2) with a Fisher exact test with R (R-Core-Team, 2018). Each test was performed within a homogeneous subsample: with BP only and with GP only for the BS/IS comparison; within BS or IS only for BP/GP comparisons. For these comparisons, we excluded uninterpretable profiles. We also tested the proportions of loci that were correctly amplified and those that presented an uninterpretable result (i.e. profiles with an obvious but fuzzy amplified product) with the same test and following the same steps as explained above.

A substantial proportion of homozygous profiles could be due to amplification failures of one allele in a heterozygous genotype (Kaboré et al., 2011), particularly so if homozygous profiles outreach the frequency of 1/*K* expected in clonal populations (see Appendix 1). Given the amplification failures observed in our study, we used a one-sided Spearman’s correlation test to test whether there is a positive relationship between the proportion of blank, of uninterpretable genotypes (among visible genotypes), of total missing profiles (blanks+uninterpretable) and the proportion of homozygous profiles (among loci with a clear genotype) found per individual samples.

To visualize the organization of individual samples among each other’s and in particular individuals from animals as compared to *Tbg*1 reference stocks, we built a Neighbor-Joining-Tree (NJTree) (Saitou and Nei, 1987) from a Cavalli-Sfoza and Edwards chord distance matrix (Cavalli-Sforza and Edwards, 1967), as recommended (Takezaki and Nei, 1996). We computed genetic distances with FreeNA (without INA correction) (Chapuis & Estoup, 2007), and constructed the tree with Mega 7 (Kumar et al., 2024). For this analysis, we kept only samples that had more than 70% of interpretable loci to avoid empty lines in the matrix and to limit biases in genetic distance estimates.

We defined five hierarchical levels of potential subdivision for this trypanosome population. The smallest level corresponded to the 24 villages sampled, contained in eight village groupings, in three foci, and in the total sampled area (see Figure 1).

Considering that population differentiation becomes significant between subsamples from the same site but separated by six months (Kaboré et al., 2011), we split subsamples separated by more than six months as belonging to different cohorts (C). Sampling extended from December 2015 to May 2017. This resulted in two different cohorts for the present study: C1 (December 2015 to May 2016) and C3 (December 2016 to May 2017). For C1, sampling corresponded to Bonon in December 2015 and Sinfra in May 2016. For C3, sampling corresponded to Sinfra in January 2017, and to Vavoua in February and March 2017 (Table 1). It resulted that the total sample could be represented by different levels of subsamples: Village-Cohort combinations, Group-Cohort combinations (ignoring villages), Focus-Cohorts, and Cohorts.

Population genetic structure was evaluated through linkage disequilibrium (LD) tests and Wright’s fixation indices (Wright, 1965). For this analysis, we kept only samples that had more than 70% of interpretable loci (i.e. 34 samples, including 2 BS and 32 IS). LD tests were performed by 10,000 randomizations of genotypes for each locus pair with the *G*-based test, which is the most powerful for combining tests across subsamples (De Meeûs et al., 2009). We undertook these tests with Fstat 2.9.4 (Goudet, 2003) an updated version of Fstat 1.2 (Goudet, 1995).

For a hierarchy with three levels (individuals in subsample in total sample), three *F*-statistics can be defined: *F*_IS_, which measures inbreeding of individuals relative to inbreeding of subsamples; *F*_ST_, which measures inbreeding of subsamples relative to total inbreeding; and *F*_IT_, which measures inbreeding of individuals relative to total inbreeding. Deviations of genotypic proportions expected under local panmixia are measured by *F*_IS_, while *F*_ST_ measures the effect of subdivision (degree of genetic isolation between subsamples) and *F*_IT_ reflects the combination of both (e.g. De Meeûs et al., 2007). Under the null hypothesis (panmixia and no subdivision), all these statistics are expected to be null. Otherwise, *F*_IS_ and *F*_IT_ can vary from −1 (one heterozygous class) to +1 (all individuals homozygous) and *F*_ST_ from 0 (all subsamples share the same allele frequencies) to +1 (all subsamples fixed for one or the other allele).

*F*-statistics were estimated with Weir and Cockerham’s unbiased estimators (Weir and Cockerham, 1984) with Fstat 2.9.4: *f* for *F*_IS_, *θ* for *F*_ST_, and *F* for *F*_IT_. These estimators vary similarly to their corresponding parameters except *θ* that can display negative values when subsample share allele frequencies that are more similar than expected under random subsampling in the same population.

Significance of deviation from random mating or of subdivision was assessed with 10,000 permutations of alleles between individuals of the same subsample (for local panmixia), of individuals between subsamples (for subdivision) and of alleles between individuals over all subsamples (for global panmixia). The statistics used for these randomization tests were Weir and Cockerham’s unbiased estimators of *F*_IS_ and *F*_IT_ (local and global panmixia respectively) (Goudet, 1995) and the *G* statistics for subdivision (Goudet et al., 1996) computed over loci, which is the most powerful combination procedure (De Meeûs et al., 2009). All these tests were undertaken with Fstat.

Confidence intervals at 95% (95%CI) of *F*-statistics were computed with the Jackknife standard error over subsamples or loci; or through 5000 bootstraps over loci. These were computed with Fstat. More details can be found elsewhere (De Meeûs et al., 2007). Nevertheless, when the number of subsamples was too low, we used the variation across alleles to compute the Student’s *t* based 95%CI of *F*-statistics for each locus.

According to Goudet et al.’s criterion (Goudet et al., 1994), the level at which hierarchically nested subsamples are significantly subdivided can be assessed by comparing their *F*_IS_ (De Meeûs et al., 2006). Comparisons between *F*_IS_ obtained within village-cohort combinations (*F*_IS-villages_), within groups (ignoring villages) (*F*_IS-groups_), within foci (ignoring villages and groups) (*F*_IS-foci_), within each cohort (ignoring villages, groups and foci) (*F*_IS-cohorts_) and within a single sample regrouping all individuals (*F*_IS-all_) were undertaken with a one-sided Wilcoxon signed rank test for paired data with Rcmdr, the pairing criterion being the locus. The alternative hypothesis was *F*_IS-villages_<*F*_IS-groups_<*F*_IS-focis_<*F*_IS-cohorts_<*F*_IS-all_, e.g. meaning that village is a better suited subdivision, because ignoring those into groups produces a significant increase of *F*_IS_ (Wahlund effect) (and so on). We also compared *F*_ST_, *F*_IT_ and the proportion of locus pairs in significant LD (%SigLD). For *F*_ST_, the Wilcoxon test was also one-sided, but in the opposite direction (Brito et al., 2025). For *F*_IT_ comparisons, we undertook a two-sided Wilcoxon test, as we could not predict a clear effect of a Wahlund effect on that statistic. The different %SigLDs were compared with Fisher’s exact tests with Rcmdr, with a two-sided alternative hypothesis, as Wahlund effects can produce a decrease of LD in populations with strong linkage disequilibria, in particular in clonal populations (Prugnolle and De Meeûs, 2010; Manangwa et al., 2019). Finally, the effect of regrouping all individuals into one sample (all) was tested for *F*_IS_ and %SigLD only.

Following theoretical results from the literature, *F*_IS_ is expected to display negative values in clones (Balloux et al., 2003), with a relatively small variance across loci (De Meeûs et al., 2006). We also expect that the variation across loci be explained by the level of polymorphism of these loci (e.g. through different mutation rates) (Séré et al., 2014). Indeed, in clonal populations with an infinite allele model of mutation (IAM), *F*_IS_=-(1-*H*_S_)/*H*_S_ (Séré et al., 2014) . In pure clones then, a positive correlation is expected between *F*_IS_ and *H*_S_. This correlation was tested (unilateral test) with a Spearman’s rank correlation test under Rcmdr. In strongly subdivided and 100% clonal populations, other parameter values are expected: *F*_ST_=-*F*_IS_/(1-*F*_IS_) (that we labelled *F*_ST-C_) and *F*_IT_=0, which can never occur with any amount of sex, unless there is full panmixia (i.e. all *F*-statistics are null) (De Meeûs and Balloux, 2005) . We thus compared *F*_ST_ and its 95%CI to those of *F*_ST-C_, and checked their equality by a Wilcoxon signed rank test for data paired by loci. We also checked that *F*_IT_ and its 95%CI did not significantly deviate from 0, and confirmed this with the permutation test described above. When variation of *F*_IS_ across loci is strong and not explained by *H*_S_, it was suggested that it may come from either very rare sex or presence of amplification problems (e.g. null alleles, short allele dominance or stuttering) (Balloux et al., 2003; De Meeûs et al., 2006; Séré et al., 2014). There is a way to discriminate between the two causes (rare sex versus amplification problems) (Séré et al., 2014). For this, we needed to compute the difference between *F*_ISexp_=-(1-*H*_S_)/*H*_S_ and the observed one (*F*_IS_obs_): *ΔF*_IS_=*F*_IS___exp_-*F*_IS_obs_. When |*Δ*FIS|≤0.05×|*F*_IS_exp_| the two values are considered superimposed, i.e. total clonality without amplification problems. When superimposed points proportion across loci *p*_sup_≤30%, rare sex better explains the data, while *p*_sup_>80% reflects more amplification problems with total clonality or negligible amounts of sexual recombination (i.e. clonal rate *c*≥0.9999). This criterion, that we will call Séré’s criterion, was applied to each locus with *H*_S_>0.5 as recommended (Séré et al., 2014).

Another expectation in strongly subdivided clones is that a large proportion of locus pairs should display a significant LD, and that this proportion does not increase, or even decrease with Wahlund effects (Prugnolle and De Meeûs, 2010; Manangwa et al., 2019).

In clonal populations, a strong correlation is expected between individual-based genetic distances of independent sets of markers (Tibayrenc et al., 1990; Telleria et al., 2013). We studied the match between Cavalli-Sforza and Edwards chord distance matrices obtained with BP and GP with a Mantel test. Matrices were obtained as described above and Mantel test undertaken with Fstat with 10,000 permutations.

A more detailed explanation of theoretical expectations in clonal populations can be found in Appendix 1.

We tried using several recent programs specific to clonal populations: MLGSim (Stenberg et al., 2003) and GenAPoPop (Stoeckel et al., 2024).

Isolation by distance test was undertaken with Rousset’s approach (Rousset, 1997) where, in a two dimension framework, Rousset’s Index *F*_ST_R_=*F*_ST_/(1-*F*_ST_) follows *F*_ST_R_=*a*+*b*×Ln(*D*_G_), and where *a* is a constant (intercept), Ln(*D*_G_) is the natural logarithm of the geographic distance between two sites and *b* is the slope of the regression. The geographic distance was computed from GPS coordinates in decimal degrees with the package geosphere (Hijmans et al., 2019) of R (command distGeo). Because clonal reproduction tends to reduce *F*_ST_ values (Balloux et a*l*., 2003; De Meeûs and Balloux, 2005; De Meeûs et al., 2006), Rousset’s model cannot be used to estimate demographic parameters from the slope of isolation by distance regression. Since some subsamples are not contemporaneous, we undertook the regression with distances between contemporaneous subsamples only (i.e. same cohort). Weir and Cockerham estimator of *F*_ST_ and bootstrap (5,000) confidence intervals were computed between each contemporaneous subsample pairs with FreeNA (Chapuis and Estoup, 2007) (without ENA correction) and then transformed into *F*_ST_R_ and regressed against Ln(*D*_G_). Isolation by distance was considered significant if the 95%CI of the slope did not contain 0. In case of no significance, we also undertook a Mantel test with Fstat with the Cavalli-Sforza and Edward’s chord distance matrix (Cavalli-Sforza and Edwards, 1967) and 10,000 randomizations because this test can be more powerful in some instances (Séré et al., 2017). This was done to avoid type II error, which is expected to be frequent for these tests (Séré et al., 2014). We converted the bilateral *p*-value of Fstat into a unilateral one (correlation was expected to be positive) as described in Manangwa et al. (2019): in case of positive correlation, the *p*-value of Fstat was halved, otherwise the unilateral *p*-value was 1-half the bilateral one. For subsample sizes issues, this analysis was undertaken for C3 in Sinfra and Vavoua only.

Due to small subsample sizes and missing data, when not enough complete genotypes were available and Fstat could not undertake the subdivision test, we used the *F*_ST-RH_ (Robertson and Hill, 1984) based permutation test of Genetix (Belkhir et al., 2004) . This test has the same properties as the *G*-based one (Goudet et al., 1996) .

### Repeated tests

All paired tests produced series of tests with some degree of dependency. This would suggest using Benjamini and Yekutieli (BY) (Benjamini and Yekutieli, 2001) adjustment. Nevertheless, after reading the relevant literature more thoroughly, it appears that BY is highly conservative, and maybe prohibitively so (Reiner et al., 2003), especially for tests on proportions (De Meeûs, 2014). Moreover, for positively or moderately negatively dependent test statistics, BY presents no real FDR (False Discovery Rate) advantages as compared to Benjamini and Hochberg’s method (Benjamini and Hochberg, 2000) (BH), which is then better to be used instead (Benjamini and Yekutieli, 2001; Reiner et al., 2003). Negatively dependent tests occur when the strength of H1 (with a better chance of small *p*-values) of some tests in the series implies that other tests in the series will tend to have also small *p*-values, while other sets of tests are undertaken under H0 (with an expected uniform distribution of *p*-values centered around 0.5). Typically, in a post-hoc multiple testing after an analysis of variance or after multiple average comparisons, we expect such moderate negatively correlated *p*-values. Nevertheless, in some instances, such correlation may be strong enough to violate the assumptions needed to apply BH. This would be the case, for population genetics analyses, when different sets of loci are in strong LD (due to selection and/or physical linkage) and others are not. This is also what would be observed during differentiation tests between different pairs of subsamples taken from a population under isolation by distance, with a lot of pairs highly significant because from remote sites, and other pairs not so different (only by chance) because computed between close by sites. In such instances, detecting the true significant tests would require the use of BY. In all other cases, BH is expected to give robust and more powerful results. This is why, contrarily to what we usually undertook in previous population genetics articles undertaken by one of us (e.g. Manangwa et al., 2019), we used BH for all multiple testing situations in the present paper.

For independent tests that needed being combined, we used the generalized binomial procedure (Teriokhin et al., 2007), computed with MultiTest (De Meeûs et al., 2009), setting the number of optimal test *k*’ at *k* when the number of combined tests are *k*<4 and *k*’=*k*/2 otherwise, as advised (De Meeûs, 2014).

### Simulations

In order to check if some parameter estimates were valid, we undertook some simulations to explore the different parameters that may lead to the pattern observed in animal *Tb* population sampled in Côte d’Ivoire. We were particularly interested in the observed *F*_IS_, the number of repeated genotypes, the shape of the NJTree, and the goodness of fit and/or direction of deviation from De Meeûs and Balloux’s and Séré et al’s criteria (see above). Simulations were undertaken with Easypop v. 2.0.1 (Balloux, 2001). Different parameter sets were explored until we obtained similar results to those observed for animal trypanosome data. All simulations started with maximum diversity and ran for 10,000 generations, with 18 loci as in the trypanosome data set. We used the population genetics analyses results to help limit the possible parameter sets (see below). At the end of each simulation, we sampled 24 subpopulations of 11 individuals (as the number of visible genotypes in Gozi 2017, in the real data set).

## Results

The raw data of microsatellite genotyping and TgsGP PCR results are given for each BS (n = 60), IS (n = 40) and reference stocks (n = 27) (127 DNA in total) in the Supplementary file S1. Among the reference stocks, only the *Tbg*1 stocks were tested positive for the TgsGP PCR excepted P26F isolated from a pig in Cameroon.

### Amplification failures

Comparisons of amplification success are presented in Figure 2 and 3. It can be seen that globally, BS provided significantly more amplification failures than IS and that BP offered better results than GP. The higher proportion of amplification successes were obtained with IS genotyped with BP.

**Figure 2:**
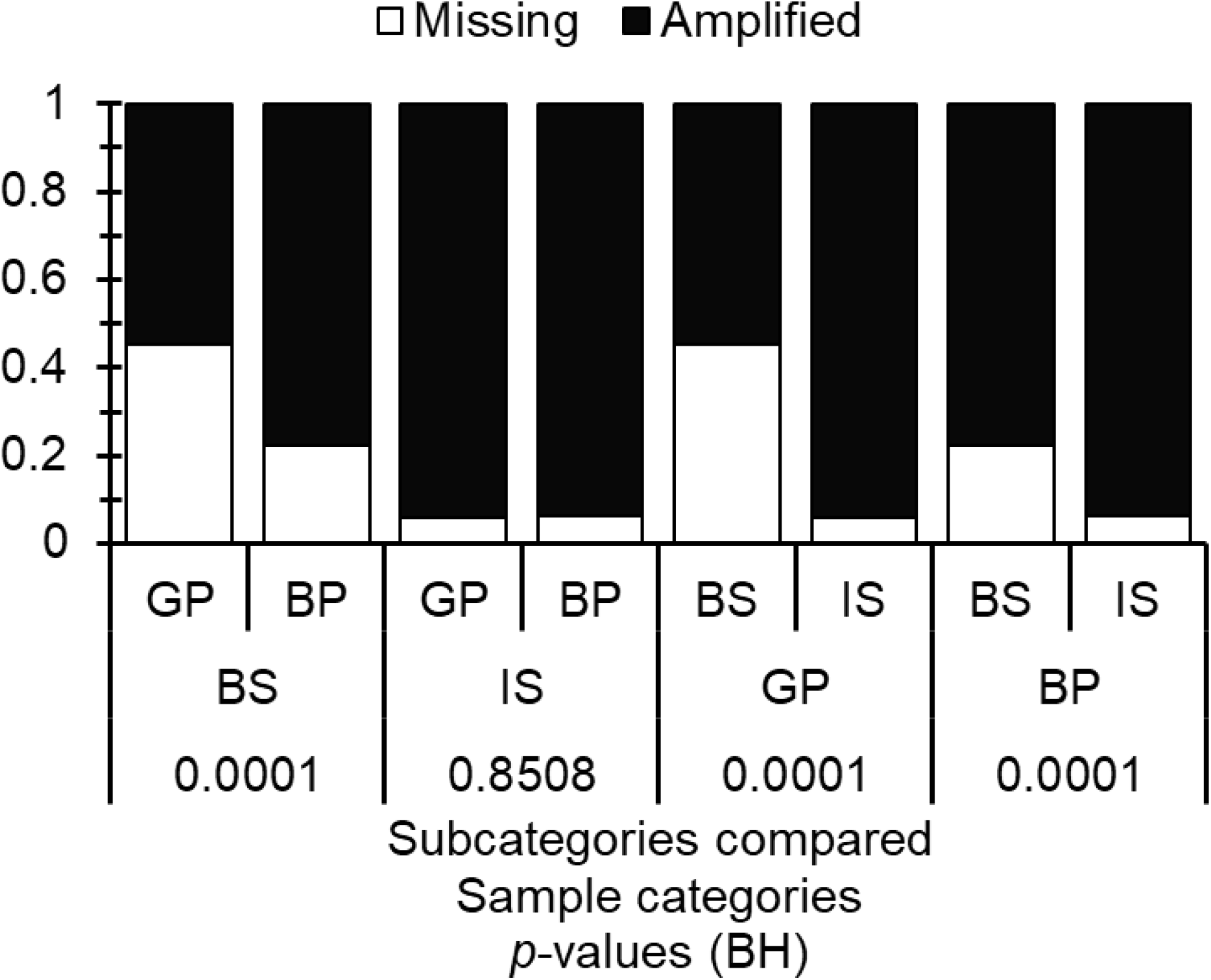
Proportions of correct amplifications (uninterpretable profiles ignored) between *gambiense* primers (GP) and *brucei* primers (BP) within biological (BS) or isolated stock (IS) samples and BS and IS within GP or BP. The different *p-values* presented were corrected with the Benjamini and Hochberg procedure for non-independent test series (BH).

**Figure 3:**
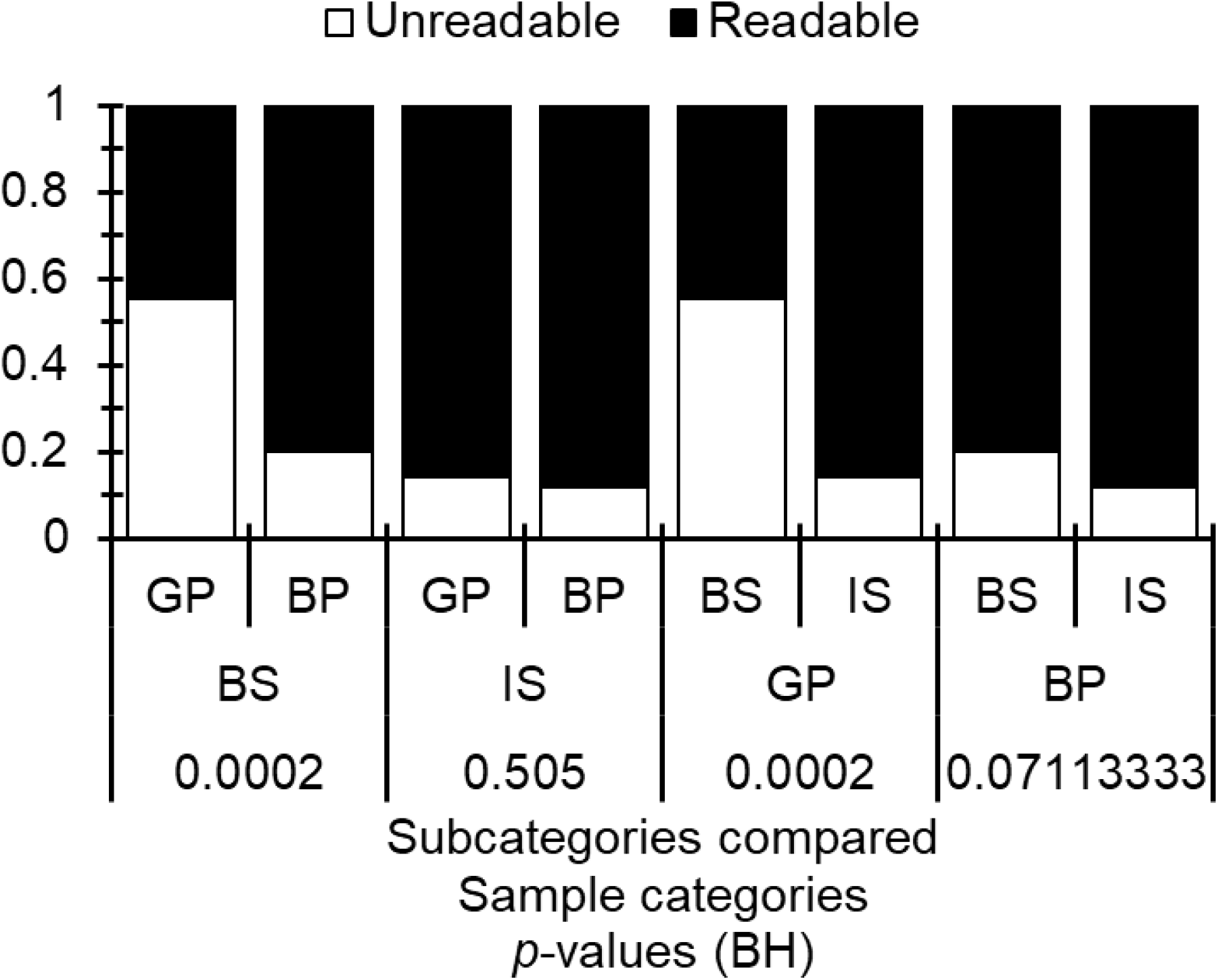
Proportions of readable amplifications (missing data ignored) between *gambiense* primers (GP) and *brucei* primers (BP) within biological (BS) or isolated stock (IS) samples and between BS and IS within GP and BP. The different *p*-values presented were corrected with the Benjamini and Hochberg procedure for non-independent test series (BH).

When focusing on reference stocks only, we noticed that *Tbb* isolates were more easily amplified than other isolates, but the difference was weak (*p*-value=0.0333). To this respect, GP did not perform better than BP (*p*-value=0.1864).

The Spearman’s correlation test showed that there was a strong and significant correlation between the proportion of blank genotypes found per locus and the proportion of uninterpretable profiles (*ρ*=0.6671, *p*-value=<0.0001). We also found a significant correlation between the proportion of blanks and the proportion of homozygous profiles (*ρ*=0.4314, *p*-value<0.0001), and between the proportion of uninterpretable profiles and homozygous ones (*ρ*=0.1762, *p*-value=0.0421). This means that the number of missing data is a fair measure of the quality of the DNA of the concerned individual: amplification failures increase the risk of uninterpretable results and the risk of dropouts of one of the two alleles in heterozygous profiles. This may also mean an underestimation of *F*_IS_. Nevertheless, when keeping stocks with more than 70% of interpretable profiles, these correlations became non-significant (all *p*-values>0.23) and even negative between blank and homozygous profiles. However, in this subset of data, the correlation between uninterpretable and blank genotypes stayed strong and significant (*ρ*=0.4065, *p*-value=0.00085). Consequently, although the last correlation confirms the lowest quality of individuals with more blank and uninterpretable genotypes, this probably did not significantly alter the estimate of heterozygote frequencies when keeping stocks with the highest amplification performances, as the one used in analyses described below.

### NJTree analysis

Out of the 100 DNA samples from BS and IS, 66 displayed more than 30% amplification failures. Therefore, only 34 samples and the 27 reference stocks were considered for this analysis. The NJTree obtained with these samples is presented in Figure 4. All *Tbg*1 reference stocks took place in a distinct clade. All other samples including *Tbg*2 and *Tbb* reference stocks and both BS and IS, occurred in a scattered order in the tree. Surprisingly, the four TgsGP positive IS looked very distant from the *Tbg*1 clade. All came from the Vavoua focus of Cohort C3, and two, from the same village (Trafla-Gottron) displayed the same multilocus genotype (same clone).

**Figure 4:**
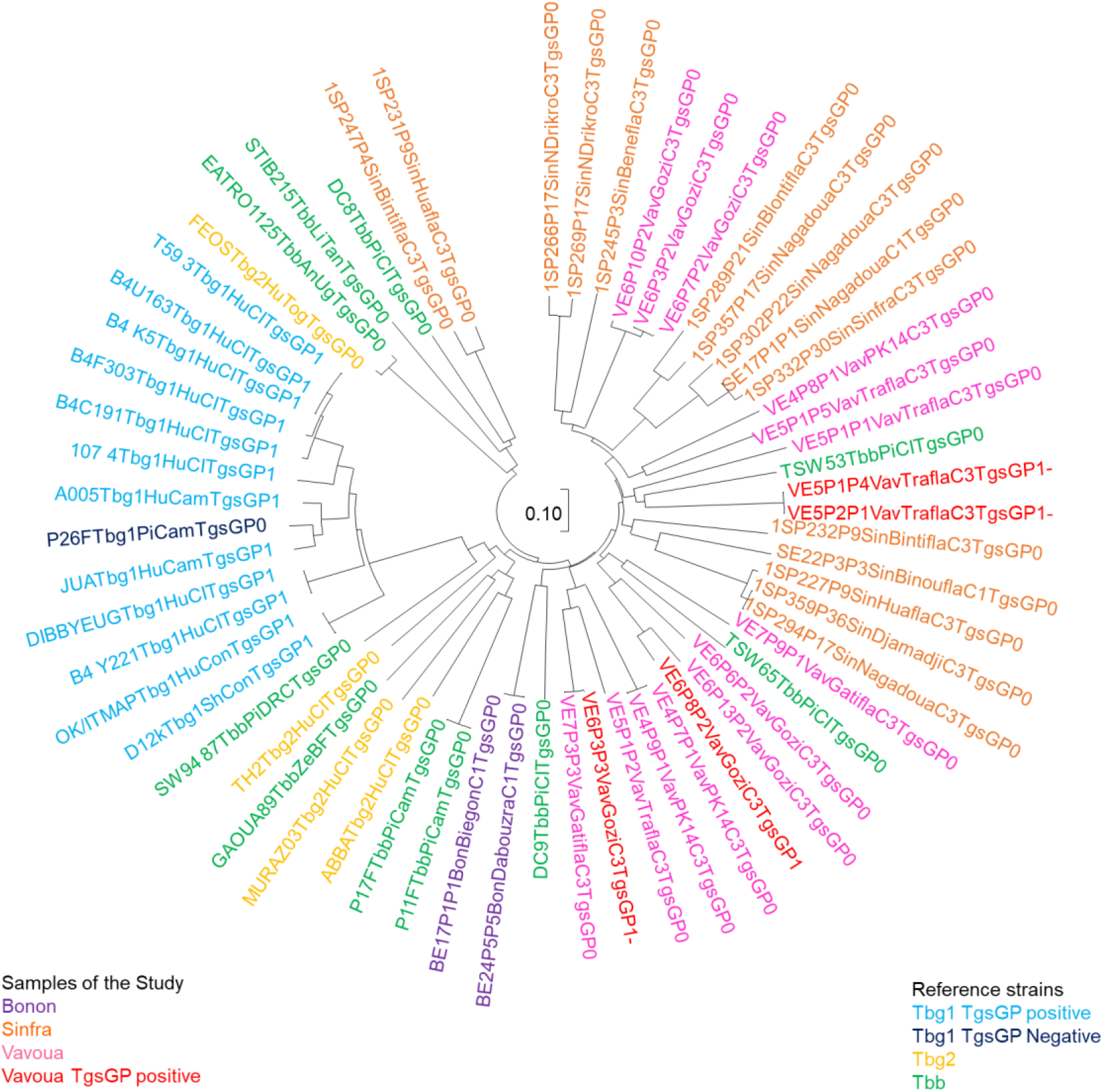
Neighbor-joining Tree (NJTree) of trypanosome stocks from a Cavalli-Sforza and Edwards chord distance matrix with the 34 individual samples that provided an interpretable profile for more than 70% of microsatellite primers among the 18 available, and 27 reference stocks. Reference stocks of *Trypanosoma brucei gambiense* group 1 (*Tbg*1), *Trypanosoma brucei gambiense* group 2 (*Tbg*2), *Trypanosoma brucei brucei*, (*Tbb*), and samples under study are in different colors, as are TgsGP positive stocks. Names of useful taxonomic units (UTOs) are explained in the Supplementary file S1 and corresponding data.

### Subdivision levels and Wahlund effects

According to the Figure 5, the most relevant subdivision level was the village-cohort combination. For sample size issues, except for cohorts and “all in one”, only cohort C3 was used for these measures.

**Figure 5:**
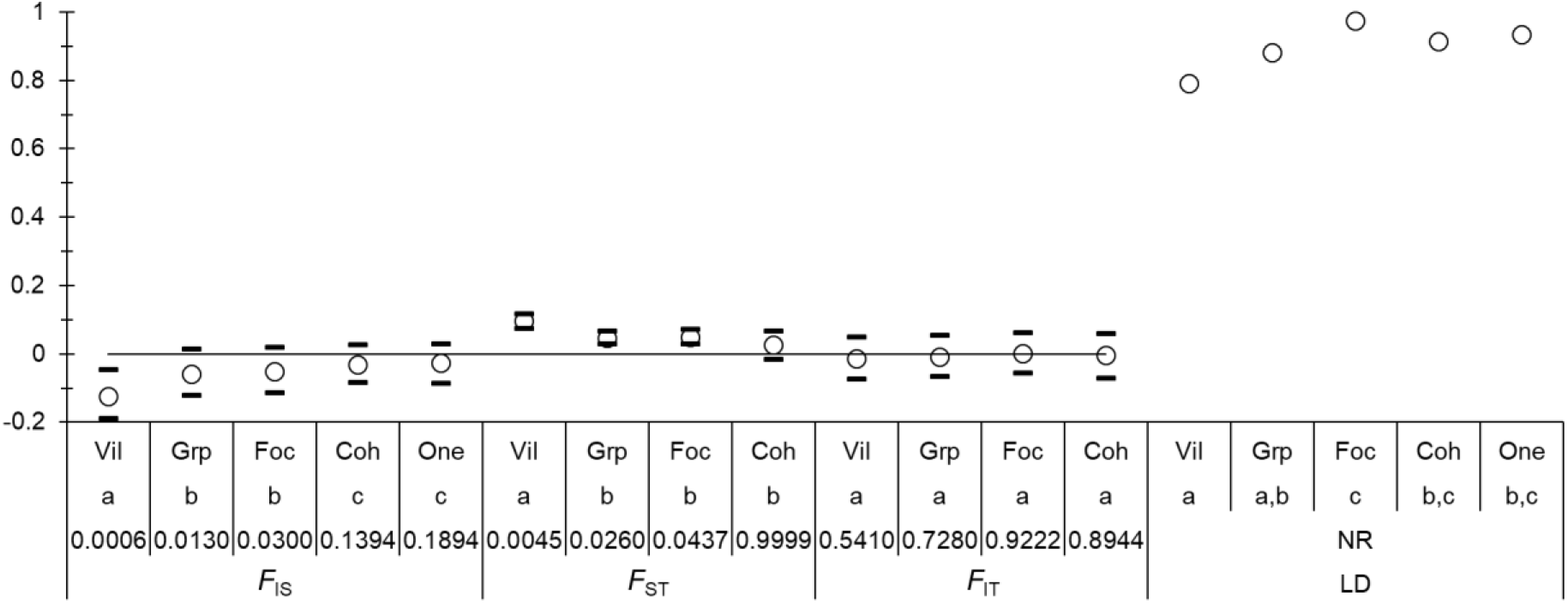
Comparisons of the different levels of potential subdivision (Vil=villages, Grp=groups, Foc=foci, Coh=cohorts, and One=all individuals grouped into one sample) as defined in the text and Figure 1, of isolates of *Trypanosoma brucei brucei* from pigs in three foci of Côte d’Ivoire, on four statistics: *F*_IS_, *F*_ST_, *F*_IT_ and proportion of pairs of loci in significant linkage disequilibrium (LD). Grouping that are significantly different are indicated by different letters (a, b or c). Significant deviation from 0 of each statistic is also indicated except for LD (NR=Not Relevant).

Consequently, we kept the village as the relevant geographic level of subdivision. Giving subsample sizes in cohort C1 we only studied cohort villages in C3 in the following.

### Evidences for clonal propagation

Within villages *F*_IS_=-0.123 in bootstraps 95%CI=[-0.189;-0.045]. It also appeared that variation across loci was very weak (Figure 6), despite the small subsample sizes and the substantial proportion of amplification failures. The correlation between *F*_IS_-village and number of missing data (blanks genotypes) was low and not significant (*ρ*=0.3463, *p*-value= 0.0796). It means that null alleles did not affect much *F*_IS_.

**Figure 6:**
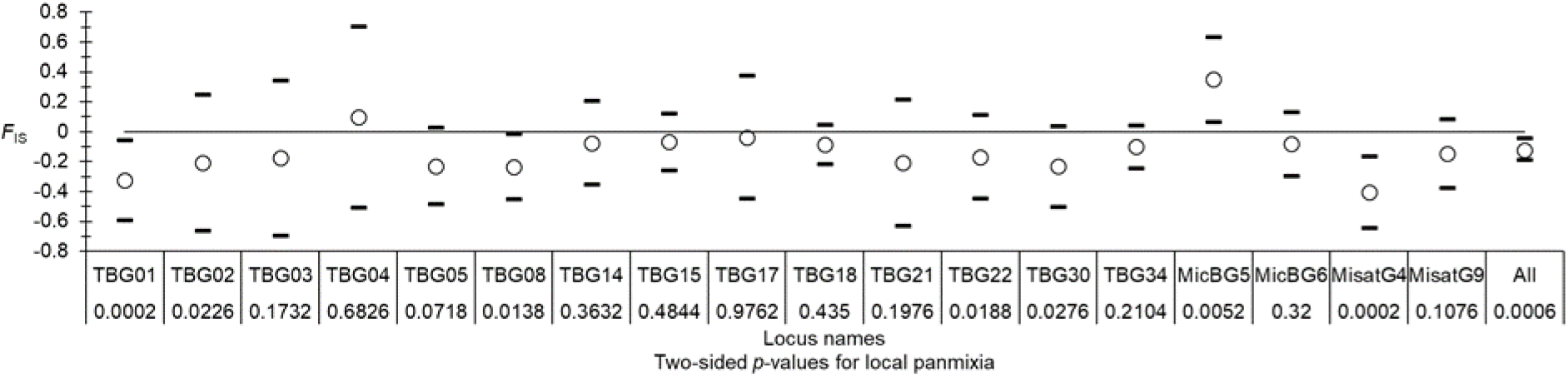
*F*_IS_ values observed in subsamples defined with cohort/village combinations (open circles) with their 95% confidence intervals (95%CI) (black dashes). The 95% CI of the average was obtained by bootstrap over loci and the 95% CI of each locus was obtained with the standard error across alleles. The two-sided *p*-values obtained after randomization of alleles between individuals in each subsample are indicated.

The correlation between *F*IS and *H*S was positive and significant (*ρ*=0.4469, *p*-value=0.0323).

Nevertheless, Séré’s criterion was never met since *p*_sup_=0%. This comes from the fact that genetic diversity, though reasonably high (*H*_S_= 0.708), appeared not high enough for the level of *F*_IS_ observed to fit Séré et al’s criterion. Said otherwise, *F*_IS_ was not enough negative, given the level of *H*_S_ observed in the dataset.

The correlation between BP and GP genetic distance matrices was positive and highly significant (*ρ*=0.23, *p*-value<0.0001), as expected in strongly clonal populations.

According to De Meeûs and Balloux’s prediction, the *F*_IS_ observed would predict a *F*_ST-C=_–*F*_IS_/(1-*F*_IS_)=0.1095 in bootstraps 95%CI=[0.0431; 0.1590], which was not significantly different from the *F*_ST_=0.0970 in 95%CI=[0.075; 0.1180] (*p*-value=0.4951). Additionally, *F*_IT_=-0.014 in bootstraps 95%CI=[-0.073; 0.05] was not significantly different from 0 according to bootstraps and randomization tests (*p*-value=0.541). In line with these results, we indeed observed a strong proportion of locus pairs in significant LD, and an evolution with Wahlund effects that appeared stable (Figure 5). These observations are in line with a total clonality in a strongly subdivided population. If so, the number of immigrants entering a subpopulation per generation could be computed as (De Meeûs and Balloux, 2005) *Nm*=–(1+*F*IS)/(4*F*IS)=1.78 in 95%CI=[1.07; 5.31]. The trypanosomes from pigs sampled in different regions of Côte d’Ivoire thus seemed to behave as a completely clonal and highly subdivided population. Given the *F*IS range, quite far from −1, subpopulations probably displayed important sizes and then very weak immigration rates.

MLGSim failed to provide any reliable result on the only subsample with enough isolates (Gozy for Cohort 3). For GenAPoPop, only Sinfra and Nagadoua (C1 and C3) could be analyzed. We used default parameters for the analysis plan. The best probabilities were rather small (close to 0 and 0.1 for Sinfra and Nagadoua respectively), with 10% of clonality and 80% of selfing in both villages. Such figures are incompatible with our observations. These results are probably entirely explained by the particular sampling we had to deal with and within a strongly subdivided population at very small scales, leading to very small subsample sizes.

### Isolation by distance within cohort 3 and subdivision

The slope of isolation by distance did not significanty excluded 0 over all subsamples (*b*=0.0117 in 95%CI=[-0.0063; 0.0239]). The Mantel test outputted a marginally not significant *p*-value=0.0526, but with a bimodal distribution of distances, all between 0 and 20 km within foci, or between 100 and 200 km between foci. Within foci, the Mantel was not significant (*p*-value=0.113). Nevertheless, subsample sizes and/or numbers being rather small, we could not exclude a type error II. For information, a one-sided paired Wilcoxon test between *F*_ST_ across all subsamples and the average (weighted with subsample sizes and genetic diversity of each locus) measured within foci only, provided a highly significant difference (*p*-value=0.0002), with *F*_ST-all_=0.097 in 95%CI=[0.075; 0.118] (*p*-value=0.0045); *F*_ST-foci_=0.0625 in 95%CI=[0.0065; 0.1127] (generalized binomial *p*-value=0.2492, combining the two *F*_ST-RH_ based tests with *k*’=2). Isolation by distance may thus be there but not visible with the sample size available.

### Simulations

For purely clonal simulations, we used expectations from *F*_IS_ to estimate *Nm*=-(1+*F*_IS_)/(4*F*_IS_) (De Meeûs and Balloux, 2005) and *N*=-(1+*F*_IS_)/(4*uF*_IS_) (Simo et al., 2010) to limit possible values for subpopulation size *N* and migration rate *m*. When sex was involved, we used *Nm*=(1-*F*_ST_)/(4*F*_ST_). We then used different values of mutation rates *u*, number of subpoplations *n*, and number of allelic states *K* that are presented in the supplementary file S2.

During the exploratory phase, any simulation with a clonal rate *c*<1 failed to produce data sets in agreement with De Meeûs and Balloux’s criterion. We then explored the case *c*=1 for various subpopulation numbers (*n* between 25 and 100), subpopulation sizes (*N* between 1000 and 10000), immigration rates (*m* between 0.001 and 0.0001), mutation rates (*u* between 10^-5^ and 10^-3^) and maximum number of alleles (*K* between 18 and 30). We then compared the shape of the NJTree obtained, the *F*_IS_, *F*_ST_ and *F*_IT_ to those observed in the real animal trypanosome data set. The best fit was obtained for *n*=25, *N*=1000, *m*=0.001, *u*=0.001 and *K*=18. No significant difference was obtained between the values observed in the real dataset and those obtained in this simulation (Figure 7 and 8).

**Figure 7:**
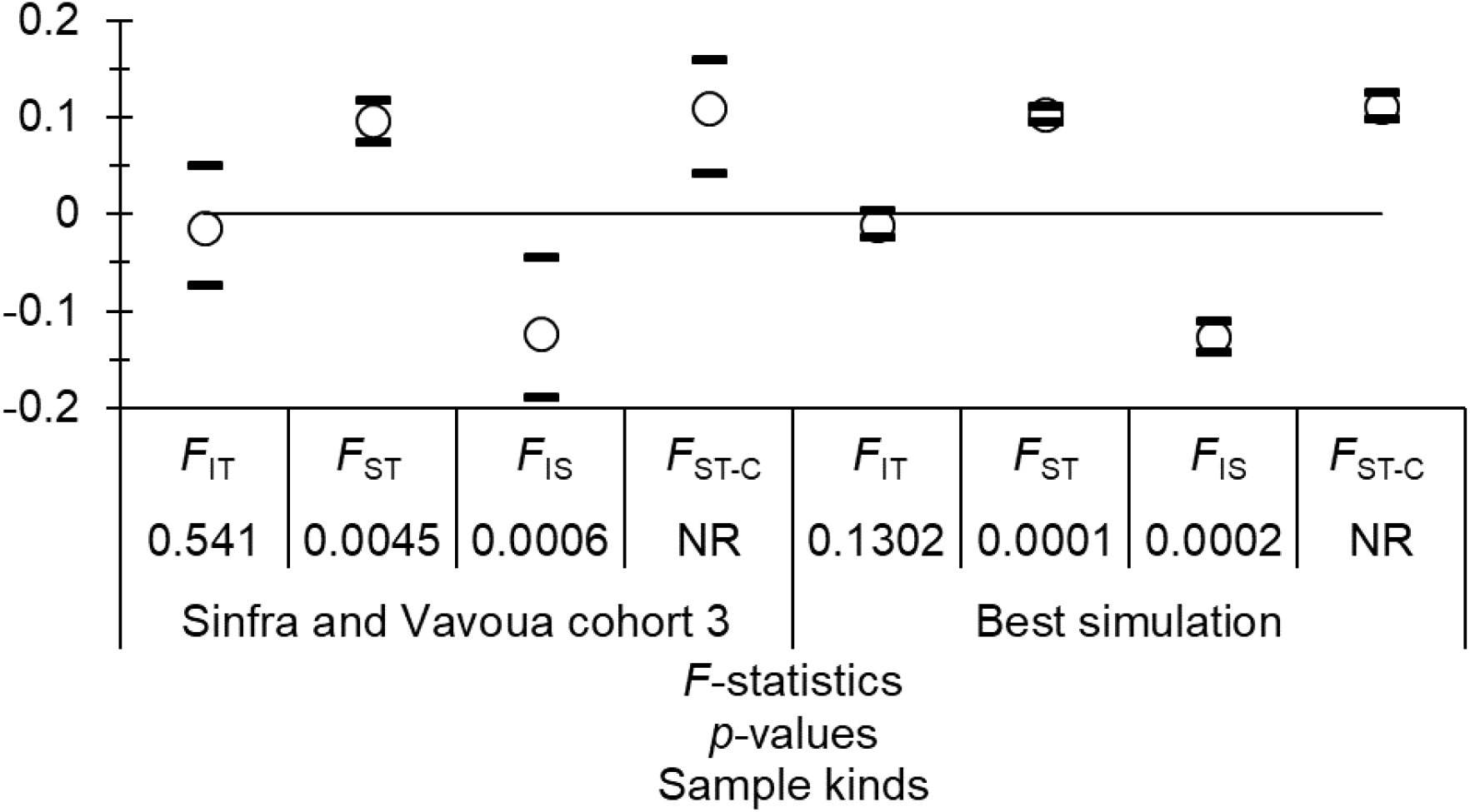
Comparison of *F*-statistics estimates (*F*_IS_, *F*_ST_ and *F*_IT_) between the real values observed in trypanosomes from Côte d’Ivoire for cohort C3 in Sinfra and Vavoua foci, the values obtained with our best simulation, and the values of *F*ST expected in case of strongly subdivided clonal populations (*F*_ST-C_=-*F*_IS_/(1-*F*_IS_)). For each parameter, 95% confidence intervals (95%CI) obtained by 5000 bootstraps over loci are represented.

**Figure 8:**
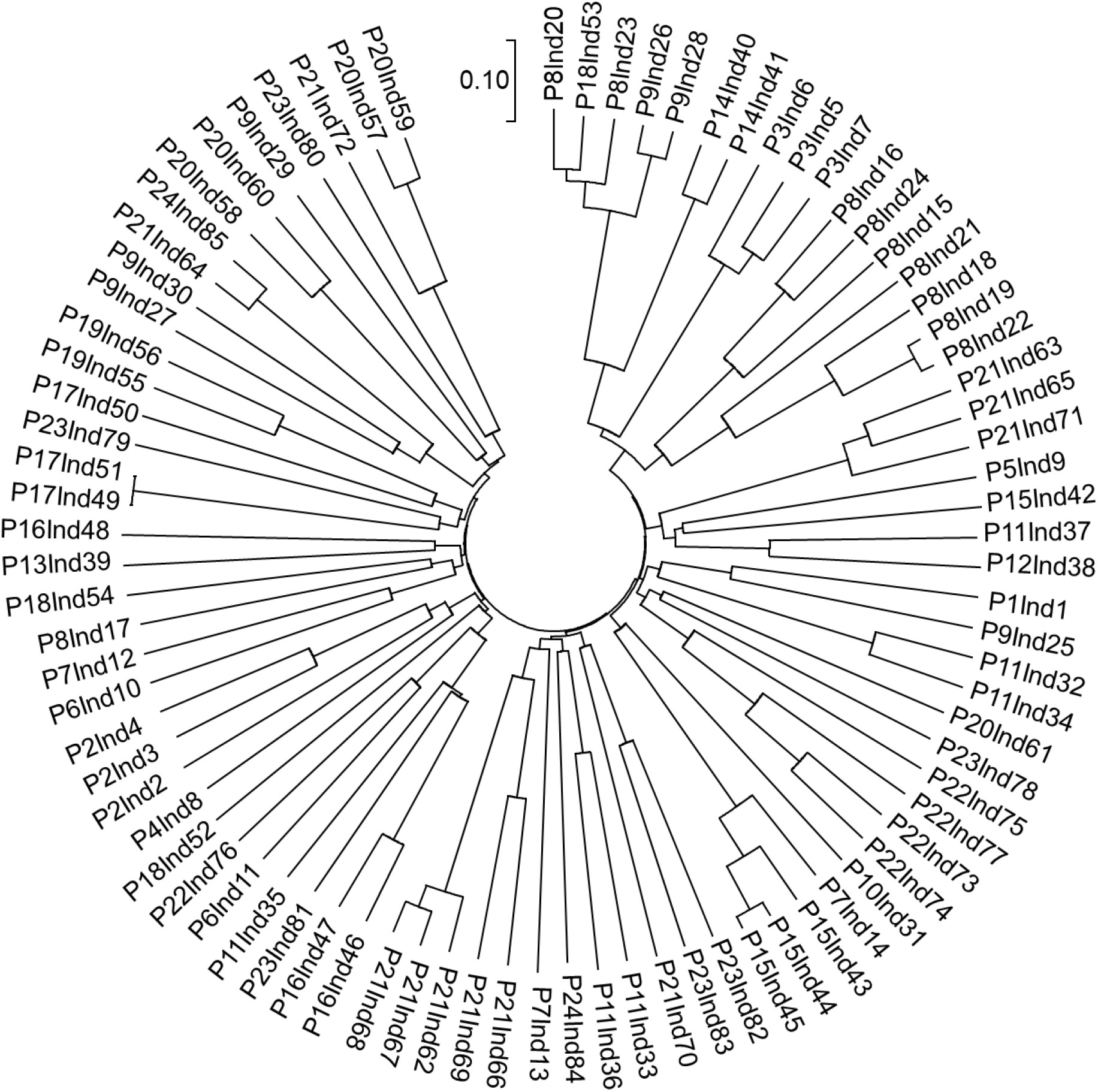
Cavalli-Sforza and Edward’s chord distance based NJTree obtained with the simulation with the best fit to animal trypanosomes subsamples from Côte d’Ivoire. Identical multilocus genotypes display a 0 distance (one instance in that case with individuals 51 and 49).

## Discussion

In Côte d’Ivoire, pigs appeared as able to play an important role in the transmission, the propagation and the spread of *Tb* s.l. parasites. In this context, the characterization of trypanosomes found in pigs could give a real indication of the genetic diversity of *Tb* s.l. circulating in Ivoirian HAT foci, clarify the relationships between *Tbb* and *Tbg* and improve our comprehension of HAT epidemiology.

Before interpreting the population genetic approach results, we have to discuss the surprising results obtained with the TgsGP and microsatellite PCR. TgsGP positive individuals found in pigs should indicated the presence of *Tbg*1 in this host, whereas it no longer seemed to circulate in humans, even though both hosts share the exact same environment as already discussed (Traoré et al., 2021). Indeed, while TgsGP is described as specific of *Tbg*1, the four TgsGP-positive IS from Vavoua did not fit into the *Tbg*1 clade according to the NJTree (Figure 4).

In a recent study targeting the TgsGP gene to detect *Tbg* in tsetse flies in Uganda, the TgsGP primers were suspected to cross-react with DNA from an “unidentified source” after sequencing the DNA products (Cunningham et al., 2020). Although the TgsGP gene is an identification target for *Tbg*1 (Gibson et al., 2010; Radwanska et al., 2002), similar genes have been identified in other *Tb* s.l. isolates (Felu et al., 2007). According to Gibson et al. (2010), TgsGP-like genes can be found in some isolates of *Tbb*, *Tbr* and *Tbg*2. These genes could harbor sequences closely related to those targeted by the TgsGP primers generating false positives as is possibly the case in the present study.

Another surprising result of the present study is the occurrence of a TgsGP-PCR negative result for a *Tbg*1 reference stock (sample P26F in Figure 4). One feature of this stock is that it was isolated from a pig in Cameroon and was used to argue for the existence of an animal reservoir of *Tbg* (Nkinin et al., 2002; Simo et al., 2005; Simo et al., 2008). Indeed, it was previously identified as *Tbg*1 in several other studies using different various methods of genetic characterization. It was also confirmed as *Tbg*1 in the present study using microsatellite loci.

These results show that there is a lack of knowledge about the taxonomy of *Tb* s.l. and the boundary and interactions between its different entities (subspecies) that can potentially share the same hosts. TgsGP and TgsGP-like genes are part of the variant surface glycoprotein genes (VSG), which allow some trypanosomatid parasites to evade the mammalian host’s immune system by extensive antigenic variation (Blundell et al., 1998). Additionally, TgsGP gene is essential in the mechanisms that confer resistance to human serum (Capewell et al., 2013). Assessing the human serum resistance of these particular stocks and studying the mechanisms involved will require further studies.

Nevertheless, our results showed that TgsGP PCR results should be interpreted with caution as they may lead to an erroneous estimation of the *Tbg* prevalence as recently observed in HAT foci in Côte d’Ivoire (Ilboudo et al., 2025).

Another limit is that both TgsGP and microsatellite primers target a single DNA sequence copy and may not be sensitive enough to detect the targeted species. Given the low parasitaemia generally observed in *Tbg* infections in human patients (Deborggraeve and Büscher, 2012), it is thus possible that *Tbg* parasites could be even rarer when in an animal host and missed by one of these two methods.

For the 18 microsatellite loci considered here, a substantial level of genetic diversity was found in *Tb* stocks circulating in animals, as illustrated by the Figure 4 and the relatively high level of genetic diversity measured. However, the primers used for this study revealed a significant amount of amplification failures during genotyping. This is particularly critical for blood samples (BS), meaning that population genetics of trypanosomes appears much easier with isolated stock sample (IS) (Figure 2 and 3). Applying primers designed from a particular species, subspecies or remote population to more or less genetically distant related taxa may generate amplification problems (De Meeûs et al., 2019). Stocks from pigs all appeared as *Tbb* and it is not surprising that *brucei* primers (BP) amplified better these stocks than primers defined from *T b gambiense* (GP). Nevertheless, there was still important numbers of amplification problems, whatever the primer kind. The BP where designed from the genomic sequences of clone TREU 927/4 (Koffi et al., 2007). Clone TREU 927/4 was derived from the isolate GPAL/KE/70/EATRO 1534 that originated from Kiboko, Kenya. It was isolated from a *Glossina pallidipes* fly in 1970. It has a degree of resistance to human serum, though it lacks the SRA gene that is characteristic of the human infective subspecies *Tbr* from East Africa (Gibson, 2012). It is thus possible that this clone is genetically very distant from the different stocks we used in the present study, hence explaining the important amplification failures met. The correlation between loci homozygosity and missing data (Kaboré et al., 2011), proves that the primers have difficulties to produce correct PCR products. This suggests that there is an actual need for the development of other markers, may be from cloned samples that will require being selected as most representative of the diversity met in all the *Tbb* stocks of the present study. The fact that some primers failed to amplify some samples confirms the results of other studies (Koffi et al., 2007; Morrison et al., 2009), especially on field samples without isolation of parasites. The low quantity of DNA for some *Tb* genotypes can also favor amplification failures and may explain the lack of some genotypes due to the presence of unreadable alleles.

These amplification failures led to exclude all samples with less than 30% of locus amplification success. With the remaining individual samples, we showed that heterozygosity was accurately estimated, and the remaining missing data due to random allelic dropouts. This thus led to a rather accurate estimation of *F*-statistics in that sub-dataset. There were convincing indications that the populations investigated here was highly or more likely absolutely clonal and strongly subdivided, according to most criteria, except for Séré et al’s superimposition criterion. This apparent contradiction may come from a higher homoplasy and mutation rate as compared to what was explored in Séré et al.(2014). Simulations suggested that the results observed in these *Tbb* individual samples were compatible with a big clonal population (*N*_total_=25,000), composed of strongly isolated demes (immigration rate *m*=0.001), and that the markers used displayed very high mutation rates (*u*=0.001) with some degree of homoplasy (number of possible alleles *K*=18). A better fit could have been reached if we had explored subpopulations of variable sizes and variable values for *u* and *K* across the 18 loci to adjust to trypanosome data, but this would have been extremely time consuming and out of the scope of the present paper. The results presented in the Figure 7, in particular, are not compatible with the existence of any detectable recombination rate (e.g. less than 1/10.000 (De Meeûs and Balloux, 2005). Absolute clonality is in contradiction with the belief that *Tbb* extensively sexually recombine in the wild (Capewell et al., 2013; Echodu et al., 2015). Comparison with other data is however difficult. First, we did not use the same genetic markers. In addition, we took care of working with contemporaneous subsamples, avoiding temporal Wahlund effects, and working at the smallest and most relevant geographical scale (villages). Date of sampling was not controlled in the studies from the cited literature, with isolates that may extend over, at least, 10 generations (Capewell et al., 2013) or even 150 generations (Echodu et al., 2015). Moreover, the parameter estimates variation across loci was much smaller in our data than in other published datasets. Additionally, genetic diversity appeared much smaller (Capewell et al., 2013) or even very weak (Echodu et al., 2015) as compared to what we observed with our microsatellite loci. This issue will require further investigations to ascertain the exact reproductive strategy of *Tbb* in the different zones where it can be found, but our results suggest that *Tbb* may be much more clonal in the wild than previously speculated. We also found that *Tbb* populations may be subdivided at scales as small as or even smaller than discs with a radius of 10 km, as defined by the villages concerned by our study.

Given the available data, it thus seemed that *Tb* s.l. is composed of several lineages that propagate clonally, with the rare occurrence of lineages that adapt to human hosts, as did *Tbg*1 10,000 years ago (Weir et al., 2016) . We used this divergent time as a standard, assuming a molecular clock for *D*_CSE_, and using maximum distances found in each group of the NJTree in Figure 4. This allowed calculating that data were compatible with the existence of clonal lineages as old as 27,000 years over all genotyped samples (*D*_CSE_=0.9), with slightly smaller divergences in Bonon and Sinfra (24,000 and 26,000 years respectively), and maximum divergences found in Vavoua. Average distances between *Tbg*2 stocks and others appeared smaller (22,000 years), but still more ancient than between *Tbg*1 and other lineages. If true, this would mean that transfer from animals recurrently, though rarely, occurs, with a very successful one (*Tbg*1), which propagated across all Western and Central Africa.

Our attempt using alternative and specific softwares dedicated to clonal populations (MLGSim, GenAPoPop) confirmed that such programs can only work on big subsamples localized in restricted geographic areas and can hardly be used for microbian populations as trypanosomes.

Isolation by distance seemed not to structure this trypanosome population, at a level detectable by our data at least. However, the weak connectivity between villages within foci, together with small subsample sizes may explain why we did not find any signature of isolation by distance. Wherever possible, further studies should include more stocks per village to gain in power for such kind of tests. Nevertheless, this may also be due to an almost complete isolation between villages, indicating a poor success of immigrant individuals.

Global studies on dispersal of the different actors involved in the epidemiology of African trypanosomoses due to *Tb* will be required to better interpret the results obtained during the present study.

In the context of HAT elimination, it is crucial to develop more accurate tools to study the epidemiological role of potential animal reservoir for *Tbg*. Controlled experimental infections of domestic animals, as recently performed with *Tbb* in pigs (Ilboudo et al., 2023), will be needed to evaluate the specificity and sensitivity of the currently available tools and their usefulness in assessing the role of animals in the maintenance and circulation of *Tbg*. These tools would also improve our knowledge on the role of a potential human residual reservoir of *Tbg* and on the mechanisms involved in tolerance to infection (Büscher et al., 2018). They would also make it possible to characterize the trypanosomes that infect dermal tissues in Human. Indeed, if recent experimental and field studies have confirmed the presence of trypanosomes belonging to *Tb* s.l. in the dermis (Capewell et al., 2016; Soumah et al., 2024), doubts remain about their identity as *Tbg*1 and their role in the epidemiology of African trypanosomoses.

To conclude, we confirm the limited efficiency of the TgsGP marker in characterizing unambiguously *Tbg*1. This is why it is currently still very difficult to assess the precise role of animal and/or human reservoirs in the epidemiology of HAT. Our results also suggest that the entire taxon *Tb* s.l. may be composed of different, sometimes highly divergent, clonal lineages, a few of which adapted to human hosts, among which *Tbg*1 appeared as the most successful one to date. Finally, in the foci studied, villages appeared to harbor very big and diverse populations of trypanosomes.

This may jeopardize the control of African trypanosomoses and should encourage integrated approaches to optimize control strategies against the various species of trypanosomes that infect humans and animals. Nevertheless, with almost no exchange between villages, propagation risks seem very limited, though prevalence in tsetse flies and dispersal of infected flies will need being evaluated to precise this risk.

## Appendices

**Appendix 1:** Special features of clonal populations.

Let *Q*_I_ be the probability to sample twice the same allele in a single individual of the total population, *Q*_S_ the probability to sample twice the same allele in two different individuals from the same sub-population, and *Q*_T_ the probability to sample twice the same allele from two different subpopulations. The parametric definition of *F*-statistics is (e.g. De Meeûs, 2018):

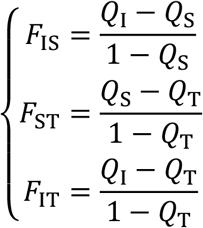

Clonality means an absence of segregation and recombination. In purely clonal populations, as a consequence of genetic drift, continuous mutation in the genome and absence of segregation, all sites tend to become heterozygous. At equilibrium, *Q*_I_ → 0. It means that *F*_IS_ → -*Q*_S_/(1-*Q*_S_), or *F*_IS_≈-(1-*H*_S_)/*H*_S_. A positive correlation is thus expected between *F*_IS_ and *H*_S_ across loci as long as the different loci display different genetic diversities (e.g. through different mutation rates *u*). This relationship allows seeing that *F*_IS_ is expected to increase with mutation rate, subpopulation sizes (*N*) and numbers (*n*) (De Meeûs et al., 2006).

According to (Séré et al. (2014), such variation across loci should lead to an equality between *F*_IS_ and −(1-*H*_S_)/*H*_S_. Nevertheless, variation across loci may also result from the presence of null alleles or rare sex. However, in purely clonal populations, null alleles are expected to have a very small impact on the *H*_S_/*F*_IS_ relationship, if rare enough (i.e. less than 20%), while very rare sex (i.e. 0.1%) will quickly completely break this relationship.

Now, if the population is strongly subdivided with a lot of possible allelic states (*K* → ∞) and the number of subpopulation big enough, we may assume that *Q*_T_ → 0. This would lead to what we call De Meeûs and Balloux’s criteria:

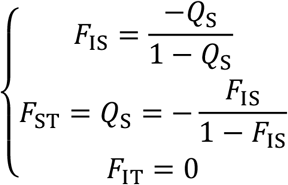

For more limited *K*’s, De Meeûs showed that, at equilibrium, *Q*_I_=1/*K* and, with limited number of subpopulations, this criterion will not hold. The criteria described above will also not hold if migration rate becomes substantial.

In clonal populations, the absence recombination generates and maintains high levels of linkage disequilibria across the whole genome. A great correlation across all markers of the genome is thus expected to occur, hence a high proportion of locus pairs in significant LD and a strong correlation between sets of markers developed independently. This absence of recombination also leads to the occurrence of identical multilocus genotypes or MLGs. In clonal populations that are strongly subdivided into big subpopulations and with important mutation rates, we thus expect the fixation of a substantial diversity of MLGs in each subpopulation, with very distant genotypes across those. Gathering a few MLGs from different and distant subsamples will not produce an increase of linkage disequilibrium (LD) between loci, as most of the time expected in “classic” sexual populations. This is why, depending on the structure of the population, Wahlund effects are expected to produce a stagnation or even a decrease in LD across locus pairs (Manangwa et al., 2019; Prugnolle and De Meeûs, 2010). This is also why, in such populations, if subsample sizes are not big enough, we cannot expect so many MLGs to emerge (see Figure 7).

## Supporting information

RebuttalLetter

## Acknowledgements

The authors thank Solenn Stoeckel and Sophie Arnaud-Haond for their useful advice for the choice of parameters for the simulations. We acknowledge all the technicians from the HAT teams of Institut Pierre Richet of Bouaké, University Jean Lorougnon Guédé of Daloa and HAT National Elimination Program of Côte d’Ivoire. We also would like to thank Fabien Halkett, Miles Anderson and one anonymous referee whose comments considerably helped to improve our manuscript.

## Author contributions

Martial Kassi N’Djetchi: genotyping, data collection, data analyses and manuscript redaction.

Thierry De Meeûs: supervision, data analyses, writing of the manuscript and design of figures.

Barkissa Mélika Traoré: data collection and manuscript correction. Sophie Ravel: genotyping and manuscript correction.

Adeline Ségard: genotyping and manuscript correction. Jacques Kaboré: supervision and manuscript correction.

Abe Allépo Innocent: data analyses and manuscript correction Dramane Kaba: supervision and manuscript correction.

Djakaridja Berté: data analyses and manuscript correction. Thomas Konan: data collection.

Bamoro Coulibaly: design of figures.

Pascal Grébaut: culture and strain isolation. Geraldine Bossard: culture and strain isolation.

Bruno Bucheton: supervision and manuscript correction.

Mathurin Koffi: supervision, data collection and manuscript correction. Vincent Jamonneau: supervision, data collection and manuscript correction.

## Data, scripts, code, and supplementary information availability

Raw data are available online as Supplementary File S1 associated with the present preprint.

