## Supplementary material for "*Trypanosoma brucei* from pigs in sleeping sickness foci from Côte d’Ivoire is structured into clonal and strongly subdivided populations": RebuttalLetter

**Rebuttal Letter**

**Revision round #1**

**Decision for round #1 : Revision needed**

Dear Recommender,

We have amended our manuscript and considered all your comments and those from the two referees. This have led to undertake almost all analyses again. During this process we realized that the stocks BE28P3P1, 1SP224P9 and VE8P1P1 displayed less than 70% of genotyped loci and were thus removed from NJTree and population genetic analyses. Alternatively, we chose to reintegrate the reference stock STIB215 (50% missing data) in the NJTree to add a Tbb representative in the tree. Members of this subspecies are indeed not frequent in the data, though extremely diverse. As could have been expected, these changes did not significantly alter our results. We also changed the data used to compare *F*-statistics between observed and simulated data. We indeed preferred comparing values observed in Cohort 3 subsamples only, in order not to include temporal effects on $F_{ST}$ and $F_{IT}$. We thus rebuilt Figure 7 (old Figure 6) accordingly.

To underline the level at which subdivision occurs, we have added a supplementary level that we labelled "group", and compared different statistics measured when considering different grouping strategies, with a new figure (Figure 5 in the new manuscript).

In the following you will find your comments and those from referees and our answer.

We must first clarify that during the revision process, we made some corrections to improve the form of the manuscript. They are visible in the track changes version, as well as those specifically related to reviewer response. However, in order to make the track changes version easier to read, we have deleted the old versions of figures and tables. We have also accepted the changes related to the bibliographic references.

We have added an affiliation in the authors list. We have also changed the order of two authors, with their consent.

The line numbers mentioned below refer to the new version of the manuscript, without track changes.

**Recommender's comments**

Dear authors,

Your manuscript has been evaluated by two reviewers. Both reviewers are experts in population genetics and clonal population genetics. Note that, despite several attempts, I have been unable to obtain an evaluation by a geneticist specialised in trypanosomes. I have also carefully read your manuscript and am providing you with some additional comments that supplement reviewers suggestions.

Both reviews found that your manuscript is an original contribution to the understanding of this Trypanosoma parasite species, responsible for the sleeping disease. The manuscript is mostly clear and the methods are sound.

On a personal note, I appreciate the transparency in the data analysis process, the authors' honesty about the amplification difficulties encountered with certain samples, and the tests and corrective measures that were used to remedy them. Both reviewers asked for some more clarifications.

Both reviewers asked for clarifications about the expectation for negative Fis value due clonal reproduction, and the variance across loci that is informative on the rate of clonality (Stoeckel & Masson 2014). I agree that the very specific case of studying the genetics of partial clonal organism populations warrants didactic explanations, especially if this manuscript will be published in the PCJ's general journal.

Answer: We have added an appendix that provide more detailed explanations on the criteria we used.

In addition to Reviewer 1’s comment, I regret that the authors did not take the usual precautions regarding the definition of clones, in relation to the observation of repeated multilocus genotypes. Admittedly, the use of 18 markers should limit the risk of accidentally observing the same combination of allele just by chance, but a formal test, such as that developed in Steinberg et al. 2003, is necessary.

Answer: We have tried using Stenberg et al (2003) MLGSim. Unfortunately, only one sample appeared big enough (11 individuals) but provided no result. The trypanosome population we are studying is subdivided into two cohorts, and into small villages that appeared to display strong genetic divergences. This imposes using quite restricted subsamples that seemed incompatible with the requirements of MLGSim. We have also added some comments on the number of MLGs in strongly subdivided but big clonal populations. As confirmed with our simulations, if we sample 11 individuals per subpopulations in such strongly subdivided clonal population, one does not expect gathering numerous repetitions of the same MLG. Accordingly, as you can see from the Figure 8 (amended manuscript), simulating such a clonal population, repeated genotypes are very rare. Nonetheless, the consistently negative $F_{IS}$, together with the very important proportion of locus pairs in significant LD, plus other criteria, confirm that the trypanosome population we are studying, probably follows this model.

We would like to make another comment here. To underline the importance of population subdivision at the smallest scale, we have added a supplementary level (Groups) and a new Figure (Figure 5 in the amended manuscript). This additional material illustrates well that these trypanosomes subpopulations are subdivided from each other's at a scale smaller than 10 km at least, with highly significant *p*-values. We also insist on the fact that this subdivision is also important enough so that we cannot pool individual samples from different villages, because doing so would produce an significant Wahlund effect.

Reviewer 2 made an interesting suggestion regarding the use of new methods to deduce the clonality rate, in addition to examining Fis values, namely using ClonEstiMate (Stoeckel et al. 2017). After reading the manuscript independently, I had the same comment. One of the strengths and originality of the dataset is that it based on time samples. Sampling two cohorts is the prerequisite for using ClonEstiMate. I believe that the dataset is perfectly suited to this and that it would reinforce the inference of the strong clonality signature discussed in this manuscript.

Answer: We used GenAPoPop, which is a more elaborate and more recent version of ClonEstimate. Only two villages could be studied. But the results were completely incompatible with our data. The program indeed outputted 10% of clonality and 80%

of selfing, which could have never led to the $F_{IS}$ we have observed. Obviously, the structure of our data was not compatible with the algorithm used in such a software. We discuss this in the revised version of our manuscript.

In line with this comment, a final section of the manuscript concerns modelling work, presenting simulations that most closely match the observed data. However, there is little detail on how this “fit” was achieved. A more formal framework, such as the use of ABC rf, would certainly strengthen this inferential work.

Answer: We thank the recommender for this suggestion. After some research, we understood that the recommender referred to DIYABC (Collin et al 2021). This software appeared dedicated to sexually and biparentally reproducing populations and does not allow looking for a reproductive strategy or even a complex population structure as what we have for the trypanosomes that are subdivided into villages within three foci. After being asked for the sex ratio of our data, we decided to abandon this approach. Nevertheless, we are open to any other tip if the recommender thinks otherwise and is incline to help us with this approach.
To bring more clarity on the approach we used, we have added some comments in the Results section and provided the different sets of parameters used in our simulations in the Supplementary File S2.

Fabien Halkett
by Fabien HALKETT, 28 Jan 2026 08:35

Reviewer 1 comments
Review by Miles Anderson, 27 Nov 2025 15:40

Line 365 says “Out of the 100 DNA samples from BS and IS, 34 displayed more than 70% amplification failure.” Should this be 30% amplification error?

Answer: Yes, the referee is right and we amended this sentence accordingly.

Line 461 "Out of the 100 DNA samples from BS and IS, 66 displayed more than 30% amplification failures".

Is there an established basis for cutoffs of 70% interpretable loci, or is this cutoff only used in accordance with the description in line 214-215 to prevent empty matrix entries? Perhaps it would be advisable to address the effects of interpretable loci cutoff percentages here or in the discussion section.

Answer: Reviewer 1 is right. This prevented empty lines in the matrix. It also allowed estimating $F_{IS}$ accurately. This was specified in the manuscript on line 226-228 " For this analysis, we kept only samples that had more than 70% of interpretable loci to avoid empty lines in the matrix and to limit biases in genetic distance estimates".
When keeping samples with more than 70% of interpretable profiles, the correlations calculated became non-significant (all p-values>0.23) and even negative between blank and homozygous profiles. However, the correlation between uninterpretable and blank genotypes stayed strong and significant (ρ=0.4065, p-value=0.00085). Although the last correlation confirms the lowest quality of individuals with more blank and uninterpretable genotypes, this probably did not significantly alter the estimate of heterozygote frequencies when keeping stocks with the highest amplification performances.

Line 414, figure 5: "cohort/village combinations (black dots)" would be clearer with black dots replaced with "open circles" in the description.

Answer: The referee is right and we have amended the legend of the figure accordingly.

Although it is mentioned earlier in the paper in lines 257-259 that FIS is expected to display strongly negative values in clonal populations in reference to Balloux et al., 2003, it is stated in lines 446-448 that FIS is far from -1 due to significant population size and very weak migration. I think this relationship between the clonality and population size is not readily apparent to the reader without directly reading the Balloux paper themselves. Perhaps, it would be useful to briefly summarize this relationship for the benefit of the reader that FIS is expected to equal -1 under strict clonality, but the relationship is sensitive to population size and mutation rates, which can result in the FIS being far from -1. As microsatellites also have generally high molecular mutation rates, this might also clarify for the reader why FIS does not reach the expectation of -1 under strict clonality.

Answer: We have added an appendix that explains more in details the expected values of *F*-statistics under pure clonality. We hope it will meet Reviewer 1's satisfaction. Regarding the independence of $F_{IS}$ from migration, it only holds in an Island model of migration at migration-mutation-drift equilibrium, if mutation is small and the number of subpopulations big. Indeed, if migration is big, immigrants will not bring a different information from other subpopulations, because of genetic exchange, while small migration rates will bring different individuals but with a very small impact. This is true also in sexually reproducing metapopulations. The extreme $F_{IS}$=-1 is a particular case where all subpopulations are fixed for two alleles at a heterozygous state. This is only expected at migration-mutation-drift equilibrium in clonal populations of size 1. The bigger subpopulations are, the farther from this limit a clonal population will be, especially if mutation rate is high. Asymptotically, in very big clonal populations, $F_{IS}$ should become close to 0.

Also, the mention of weak migration makes it sound as though migration should affect the FIS measure, but in equation 11 of Malloux et al., 2003 it is explained that FIS is independent of migration. This could also be worth mentioning that FIS is independent of migration, but migration is very weak regardless and therefore unlikely to bias the F-statistics.

Answer: In Balloux et al 2003, independence from migration occurs in a special case where the number of subpopulation is big and the mutation rate much smaller than the migration rate. In other situations, this observation does not hold any more.

Overall, I think the manuscript includes interesting, original and well-supported results that contribute to the understanding of Trypanosoma.

Review by anonymous reviewer 1, 03 Jan 2026 22:43

Summary

The authors explore the population genetics of three morphologically indistinguishable subspecies of the sleeping sickness causing agent - Trypanosoma brucei. They perform a population genetic study that suggests clonal propagation in contradiction to published data. Understanding the reproductive mode of human disease agents is critical as management will depend on the prevailing reproductive mode.

General comments

This manuscript has the potential to influence our understanding of trypanosome parasites. Overall the methods are robust but the authors need to also consider recent work in which using indirect methods, like calculating F-statistics, it is important to report the single locus Fis values rather than the mean values, even if reporting the CI intervals. Moreover, the variance in Fis is also a very good indicator of the prevailing reproductive mode and would bolster the arguments in the manuscript.

Answer: Single locus $F_{IS}$ were all presented in the Figure 5 (now Figure 6 in the amended manuscript). Moreover, the variance was indeed very small, as illustrated by the narrowness of the 95%CI.

Specific comments

L40 – microsatellite primers or loci?

Answer: The referee is right and we amended the abstract accordingly.

L41 – explain what this means – they do not fit into Tbg? How do they not fit?

Answer: We have added " that positioned those far outside the lineage of this subspecies in the dendrogram" that we hope will meet Referee 2's satisfaction.

L54 – can you expand briefly on ‘targetd for elimination’?

Answer: The first sentence of the introduction was modified but the clarification requested by the reviewer was added Line 74-77 : “With 747 and 799 cases reported in 2021 and 2022 respectively, *gambiense* HAT (g-HAT) responsible for 94% of the total number of reported cases, is currently targeted for elimination of transmission (zero new cases during a period of five consecutive years) (Franco et al., 2024).”

L70-74 – maybe integrate with the opening paragraph?

Answer: We have merged this paragraph and the 2nd one with the 1st.

L183 – provide actual names of dyes and which you used as proprietary dyes have different molecular weights than some you can get from other manufacturers. Did you use 6-FAM, NED, PET, and VIC?

Answer: We have amended the manuscript and the supplementary File S1 accordingly

L198 – what do you mean by uninterpretable result? Explain.

Answer: We have added an explanation " (i.e. profiles with an obvious but fuzzy amplified product)" that we hope will meet Referee 2's satisfaction.

L200 – could a homozygous genotype not also represent a homozygous genotype rather than a genotyping error? Is the ploidy known for these organisms?

Answer: Yes, it could, because of homoplasy, with probability 1/*K*. We have added this possibility in the amended text. We have added more explanations in the Material and Methods section "Microsatellite genotyping", precising that African trypanosomes are diploid and can practice meiotic sex, though almost never (if any) for Tbg1.

L214 – I think you mean loci not primers?

Answer: We have deleted this part of the sentence to avoid misunderstandings

L258 – but see recent work by Stoeckel et al. 2021 Journal of Heredity that shows that negative isn't necessarily what is expected for Fis, rather large variance for clonality. Did you calculate the variance in Fis? It is also best to plot and report all single locus Fis values, rather than a mean Fis.

Answer: If the referee refers to " Stoeckel S, Arnaud-Haond S, Krueger-Hadfield SA (2021) The Combined Effect of Haplodiplonty and Partial Clonality on genotypic and genetic diversity in a finite mutating population (M Orive, Ed,). Journal of Heredity, 112, 78–91. https://doi.org/10.1093/jhered/esaa062", this does not concern trypanosomes, which are strictly diploid. As explained above, regarding Referee 2's general comments, single locus $F_{IS}$ were all presented in the Figure 5 (now Figure 6 in the amended manuscript). Moreover, the variance was indeed very small, as illustrated by the narrowness of the 95%CI.

L476 – population genetic not genetics

Answer: Done

L534 – consider discussing the use of temporal analyses of genotype frequencies that could be done in the future, e.g., Clonestimate.

Answer: We have extensively answered to this remark in our answers the Recommenders' comments (please, see above)

L545 – loci not primers

Answer: Done
